# Dissecting single–cell molecular spatiotemporal mobility and clustering at Focal Adhesions in polarised cells by fluorescence fluctuation spectroscopy methods

**DOI:** 10.1101/220491

**Authors:** Esther Garcia, Jorge Bernardino de la Serna

## Abstract

Quantitative fluorescence fluctuation spectroscopy from optical microscopy datasets is a very powerful tool to resolve multiple spatiotemporal cellular and subcellular processes at the molecular level. In particular, raster image correlation spectroscopy (RICS) and number and brightness analyses (N&B) yield molecular mobility and clustering dynamic information extracted from real-time cellular processes. This quantitative information can be inferred in a highly flexibly and detailed manner, i.e. 1) at the localisation level: from full-frame datasets and multiple regions of interest within; and 2) at the temporal level: not only from full-frame and multiple regions, but also intermediate temporal events. Here we build on previous research in deciphering the molecular dynamics of paxillin, a main component of focal adhesions. Cells use focal adhesions to attach to the extracellular matrix and interact with their local environment. Through focal adhesions and other adhesion structures, cells sense their local environment and respond accordingly; due to this continuous communication, these structures can be highly dynamic depending on the extracellular characteristics. By using a previously well-characterised model like paxillin, we examine powerful sensitivity characteristics and some limitations of RICS and N&B analyses. We show that cells upon contact to different surfaces show differential self-assembly dynamics in terms of molecular diffusion and oligomerisation. In addition, single-cell studies show that these dynamics change gradually following an antero-posterior gradient.

## 1. Introduction

Cells interact with the surrounding environment via chemical and mechanical signalling. These interactions determine essential cellular processes such as cell migration, cell proliferation or cell differentiation, and play key roles in physiology and disease by regulating embryonic development, the immune response or cancer progression. There are a number of cellular structures that attach to the extracellular matrix (ECM) and act as hubs for integrins, cytoskeletal proteins, signalling molecules and metalloproteases. These structures constitute a bidirectional link between the ECM, the cytoskeleton and transcription factors enabling a fast communication based on continuous feedback between the ECM and the cell (1-3). Focal adhesions (FAs) are adhesion structures that concentrate integrins, paxillin, vinculin and other adhesion molecules, and directly link to actin stress fibres (4). A significant example of the biological importance of FAs is anoikis, a form of programmed cell death induced by impaired cell attachment to the ECM, which is regulated by several FA-proteins. FAs can be very stable or very dynamic depending on the cell type, the ECM or *in vitro* substrate they attach to, and the developmental stage of the FA, but the dynamics of the whole FA might not correlate with its molecular dynamics, since a very stable macromolecular structure might need a constant inflow and outflow of molecules to keep its organisation. Several studies have addressed the spatiotemporal dynamics that occur at FAs and all agree on the heterogeneity of the dynamics between FAs in different areas, different stages of maturation and even within an individual FA (5-7). However Digman et al have provided oligomeric information regarding the dynamics of paxillin during assembly and disassembly of FAs, showing that whereas paxillin freely diffuses in the cytosol, its diffusion dynamics are mostly defined by oligomerisation processes in which there is exchange of monomers at the growing end of the FA whereas in the retracting area, disassembly occurs through aggregates of paxillin and other molecules (7, 8).

Different techniques have been developed to quantify molecular diffusion using fluorescence microscopy. Most of these techniques are diffraction-limited, like FRAP (fluorescence recovery after photobleaching), FCS (fluorescence correlation spectroscopy) or RICS (raster image correlation spectroscopy)(9-11)while others are based on single particle tracking (12). Some other techniques are not diffraction limited as STED FCS (13, 14) and STED RICS (15). RICS, as other spectroscopic techniques, relay on the fluctuations of the fluorescence signal that occur over time due to the diffusion of molecules in and out an observation volume. The diffusion coefficient can then be calculated using an autocorrelation function (10). However, RICS differs from other techniques in that the observation volumes overlap with each other in order to calculate the diffusion coefficient (Fig. S1 A and B). The study of molecular diffusion in cells by RICS offers certain advantages. On one hand, RICS is able to resolve a variety of dynamic events that range from 0.1 to 1000 μm^2^/sec in an acquisition span of seconds to minutes (depending on the viability and photo-stability of the sample), whereas other techniques only cover smaller and discrete ranges of diffusion and can be only performed for short time acquisitions (11, 16). On the other hand, a drawback of RICS is that we only extract one global diffusion from the analysis, instead of being able to identify more than one diffusion coefficient like in FRAP or single-point FCS analyses. However the statistical robustness of RICS is higher compared to f.i. point- or scanning-FCS, since the sample size (number of pixels analysed) is usually larger. A benefit to RICS is the tailored information that can be obtained post acquisition by selecting regions of interest (ROIs) in the images acquired and by focusing on particular events that occur in a specific area, and also focusing on only certain specific temporal events occurring in particular time points along the experimental acquisition.

Data acquired for RICS can potentially be processed by number and brightness (N&B) analysis in the same dataset, giving an added value to the RICS data. N&B allows the quantification of the relative number of molecules in different stages of aggregation based on the fluorescent fluctuation spectroscopy analysis (8, 17-19). The combination of both techniques is limited though by the spatiotemporal scale of the event to be resolved. N&B usually requires higher photon-counts and thus longer dwell time, so when studying fast dynamics the same acquisition strategy might not yield quantifiable values for both techniques (20).

Both, RICS and N&B have been successfully applied to characterising the molecular dynamics of different cellular processes (21-26). Moreover, molecular diffusion of FA-related molecules and particularly of paxillin clusters have been previously studied by RICS, N&B, FCS and single particle tracking, constituting a reliable standard to study diffusion dynamics (7, 8, 11, 27-30). This paper takes advantage of previous publications on the use of paxillin as a model for RICS and N&B applied studies. This report aims at finding the sensitivity of the techniques as well as understanding the limitations of a combined approach of RICS and N&B. For this purpose we will: 1) investigate and quantify the molecular assembly and dynamics of paxillin at FAs when in contact with three surfaces with known different attachment properties; 2) investigate at single cell level the molecular assembly and disassembly processes at individual FAs.

In this work, we aimed to investigate FA dynamics in the context of cellular sensitivity to different surfaces. For that, we examined the temporal dynamics of paxillin that take place during the self-assembly and disassembly of the FA. These dynamics were analysed by measuring the molecular diffusion of paxillin and its oligomerisation state at specific stages of the observed FA life span. We first investigated whether RICS is sensitive for distinguishing differential adhesion dynamics to non-coated glass, poly-L-Lysine (PLL) or gelatin. Attachment to glass or PLL occurs via interaction between the polyanionic cell surfaces and polycationic behaviour of glass and PLL, whereas in the case of fibronectin, collagen or gelatin, cell adhesion is mediated by integrins, which can lead to a more dynamic interaction with the substrate than that established with polycationic surfaces. These surfaces not only differ in the mechanisms by which cell attachment occurs, but also in the number of cell contacts and the strength of attachment, and therefore in the molecular dynamics at these contact sites (31-33). We also studied whether the temporal resolution provided by RICS allowed detecting differential assembly and disassembly dynamics of FAs at different subcellular regions and upon contact to different surfaces. Finally, we tested whether RICS and N&B can be employed simultaneously to gain further insights on the molecular mechanisms of paxillin diffusion and aggregation at FAs in the aforementioned conditions at the single-cell level.

Finally, in this report we demonstrate that RICS analyses allow the detection and discrimination of discrete diffusion dynamics of paxillin at FAs attached to different surfaces. Moreover, this technique also permits to identify differential diffusion dynamics in diverse areas of the cell, being able to detect a gradual increase in molecular diffusion from the leading edge towards central areas of the cell. We describe that N&B can be combined with RICS, providing complementary information of paxillin diffusion and its state of aggregation. Combining RICS and N&B with a single-cell based approach, allows identifying and characterising the gradual variations in the molecular dynamics, in terms of molecular diffusion and oligomeric state, that occur during FA assembly and disassembly processes within a polarised migrating cell.

## 2. Materials and methods

### 2.1 Cell lines and reagents

MDA-MB-231 human breast cancer cells were cultured at 37<u>°</u>C and 5% CO_2_ in Dulbecco’s Modified Eagle’s Medium supplemented with 10% foetal bovine serum, 2 mM L-Glutamine and antibiotics (1,000 U/ml penicillin and 0.1 mg/ml streptomycin) (all from Sigma). MDA-MB-231 cells were transfected with paxillin-EGFP, using the Neon transfection system (Invitrogen, Carlsbad, California) applying the following electroporation parameters: 1050 V pulse voltage, 20 ms pulse width, 3 pulses.

### 2.2 Sample preparation and imaging

To characterise the point spread function (PSF), purified recombinant EGFP (Biovision, Milpitas, California) was diluted to a final concentration of 20 mM in PBS (Gibco, Thermo Fisher Scientific, Waltham, Massachusetts) or 1% BSA (Sigma-Aldrich/MERCK, Darmstadt, Germany) and transferred to a glass-bottomed μwell-chambers (Ibidi, Martinsried, Germany). Cells were seeded on 35mm dishes or μwell-chambers (both from Ibidi) alone or coated with 0.01% Poly-L-Lysine or 2% Type A gelatin from porcine skin (both from Sigma-Aldrich/MERCK) 24h prior imaging. Cells were imaged in phenol-free media supplemented with 25mM Hepes (Sigma-Aldrich/MERCK), at 37<u>°</u>C and 5% CO_2_.

Time-lapse images of migrating cells were acquired in a Leica TCS SP8 inverted confocal microscope (Leica Microsystems, Manheim, Germany) fitted with a HCX PL APO 63x/1.2NA CORR CS2 water immersion objective. EGFP was excited using a pulsed (80MHz) super-continuum white light laser (WLL) at 488nm. Emission signal was collected with a photon counting (HyD) detector (500-550nm) and the pinhole was set to one Airy unit. Images of 256x256 pixels at 16-bit depth were collected using 80.4nm pixel size and 8μs pixel dwell time, for at least 200 consecutive frames (Fig. S1 B).

### 2.3 Analysis of paxillin diffusion by raster image correlation spectroscopy

Raster image correlation spectroscopy (RICS) analysis was performed using the “SimFCS 4” software (Globals Software, G-SOFT Inc., Champaign, Illinois), as described (28) (Fig. S1 A and B). To characterise the PSF, 200 frames of freely diffusing recombinant EGFP (20 mM) were continuously collected over time in order to obtain the intensity fluctuations of EGFP as a function of time. The waist of the PSF was then adjusted by measuring the autocorrelation function of the EGFP solution at room temperature and 37<u>°</u>C with a fixed diffusion rate of 90 μm^2^/s. After adjusting the waist of the PSF, we calculated the diffusion coefficient of cytosolic EGFP, cytosolic paxillin-EGFP and paxillin-EGFP at focal adhesion sites, and confirmed that the values obtained were similar to those previously described for this technique (11, 28).

RICS analysis was performed in images after carefully observing the average intensity trace of the whole set of frames. This was done in order to spot possible photobleaching effects and avoid potential errors in the quantification of the diffusion coefficient. A dataset was discarded when the investigated set of frames showed an average intensity trace that abruptly decayed at the beginning of the collection, or when the investigated event did not show a clear disassembly process. This strategy allowed us to avoid using detrending algorithms. RICS analysis was performed using a moving average (background subtraction) of 10 to discard possible artefacts due to cellular motion and slow-moving particles passing through. The average of a range of consecutive images is subtracted from the image in the middle of this range. The moving average acts as a long band path spatial filter. Different values were used (4 and 16) and no apparent differences were observed. The autocorrelation 2D map was then fitted to obtain a surface map that, in most cases, was represented as a 2D projection with the residuals on top; using the previously characterised waist value from the PSF, and the appropriate acquisition values for line time and pixel time. For different regions of interest (ROI) analyses within the same cell, the corresponding region was drawn employing either a squared region with 64x64 or 32x32 pixel size. RICS analyses of these regions were performed as described above. The analysis of the independent sub self-assembly processes along the full acquisition time was performed as above. But in this case the selected frames (time scale) at which the even occurred was obtained from the study of the full average intensity fluctuation trace. Before performing the fitting, the new average intensity trace obtained from the set of frames that described the assembly, disassembly or maturation was carefully perused. As a general rule, we avoided abrupt intensity decays or increases; we focused in those intensity fluctuation events in which the intensity changes were following short increasing or decreasing steps.

### 2.4 Paxillin aggregation analysis by number and brightness

N&B analysis was performed as described (19, 25, 27) (Fig. S1 C and D). We used the same set of images and acquisition conditions described in section 2.3 for RICS analysis. For the brightness map, we did not use any detrending algorithm either, which has been described to be detrimental (26). As shown in Fig. S1 D and S2, from the brightness analysis we obtained an average apparent brightness map and an apparent brightness versus intensity scatter plot. We used this scatter plot to render a mirrored image of the average brightness map, where we could identify the oligomer distribution over a grey-scaled intensity map. We selected different regions in form of 255 pixel large boxes (with various widths). These selected regions were colour-coded in a similar fashion to the average apparent brightness map so we could obtain the mirrored graphs. Values of the apparent brightness that described each coloured box and the number of pixels enclosed in them, yielded the numbers that later were used to quantify the true brightness and the molecular brightness. These values were then plotted either as a normalised histogram (as shown in Fig. 1), or as a frequency histogram (Figs. 3 and 4). Taking advantage of the free molecular diffusion of the monomeric paxillin in the cytoplasm (7), we could obtain the apparent molecular brightness from these regions and use it as a reference for the analysis of the higher oligomerisation states. As in the papers mentioned above, the apparent brightness was transformed to true brightness and the molecular brightness (cpms) subsequently calculated. This approach was used for the rest of the selected regions, where we again converted the apparent brightness to molecular brightness. Then the values were used to identify multiples of the reference (monomer value). These multiples were categorised in three groups (as shown in Figs. 1, 3 and 4): monomers/background, dimer-trimers and multimers (4-12 oligomers). The analyses of ROIs were performed, similarly to those for RICS: the desired region was drawn employing either a squared region with 64x64 or 32x32 pixel sizes. The subsequent N&B analysis was performed as described above. As performed for RICS, the analysis of the independent sub self-assembly processes along the full acquisition time, was performed by the selecting the appropriate based on the full average intensity fluctuation trace and choosing the exact same frame range selected for RICS analysis.

**Figure 1:**
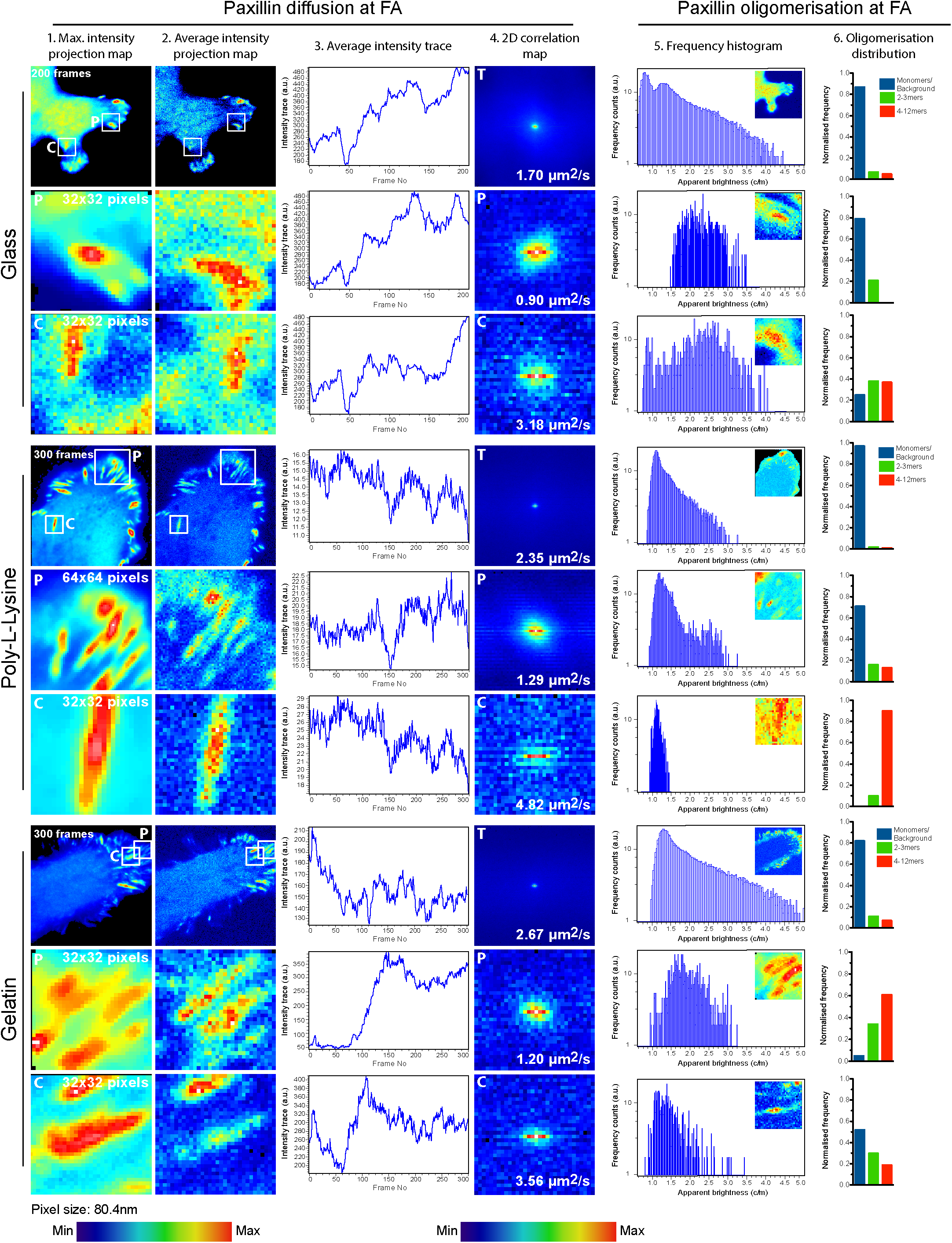
RICS and N&B analyses show distinct behaviours of Paxillin-EGFP at focal adhesions in cells grown on difference surfaces. MDA-MB-231 cells grown for 24h on glass, Poly-L-Lysine or gelatin, and imaged at 37<u>°</u>C and 5% CO_2_. RICS and N&B analysis were performed for total paxillin-EGFP (T), paxillin-EGFP at peripheral FA (P) and at central FA (C). Analyses were performed over 200-300 frames (7-10 min). Representative maximum intensity projection maps (1), average intensity projection maps (2), average intensity trace (3), 2D correlation maps and diffusion coefficient (4), resultant from RICS analysis are shown for each condition, frequency histogram (5) as well as oligomerisation distribution (6) are shown for each condition. Pixel size and colour-scale are shown below.

## 3. Results and Discussion

### 3.1 Combined RICS-N&B studies show distinct paxillin dynamics and clustering at focal adhesions in response to sensing of different surfaces

Epithelial cells attach well on glass, but they adhere more on gelatin and even more so to PLL (32, 33). In order to examine the sensitivity of the technique, we tested whether RICS is sensitive to reveal different molecular mobilities at the FA when cells sense different surfaces. To this end, we measured the diffusion of paxillin-EGFP in cells grown on different substrates: 1) non-coated glass, 2) on 0.01% poly-L-lysine (PLL) and 3) on 2% type-A porcine gelatin (Fig. 1). On average, cells attached to PLL or gelatin showed about a 30-60% faster mobility than those attached to glass, as shown in the 2D correlation maps (more elongated towards the X-axis in PLL and gelatin samples) and the diffusion coefficients shown in Fig. 1 - column 4. In addition to measuring the diffusion in a full frame over ≈7-10 minutes, we drew ROIs in specific areas of the cell, particularly in zones containing mostly paxillin located to FAs at the cell periphery and FAs further away from the leading edge of the cell (sub-peripheral or central FA from now on). Interestingly, FAs in these two regions behaved differently, as central FAs showed faster diffusion (3-4 times) than peripheral FAs (Fig. 1, column 4). Whereas peripheral FAs showed nearly no differences in diffusion among the different surfaces (ranging from 0.9-1.3 μm^2^/s), central FAs on glass presented the slowest diffusion coefficient, being the paxillin mobility of the FA in contact with PLL the fastest (Fig. 1, column 4). Therefore, RICS is sensitive not only to discern differential molecular mobility at the FA upon contact to different surfaces, but also distinguishes between adhesions at different cellular regions. This explains the discrepancies in the results obtained from regions closer to the actual size of the FA (smaller ROIs) and larger cellular regions (full-frame images) that contain several FA.

To better understand these observations, we performed N&B analyses in the same datasets (therefore same acquisition conditions) as for RICS. By these means, we could correlate information inferred from both types of analysis in spite of risking missing resolution/accuracy, though in principle, RICS and N&B analyses require slightly different acquisition parameters (19, 27). With this strategy, the N&B analysis revealed a complex oligomerisation self-assembly at every condition investigated. We could not detect differences in the frequency histograms of the apparent brightness when observing large regions of the cell (Fig. 1 column 5). However, a more detailed quantification showed some differences in paxillin oligomerisation at the FA (Fig 1 column 6; data represented in this column was inferred from the brightness vs intensity scatter plots, and the oligomerisation map shown in Fig. S2.). Quantification of oligomers by N&B analysis can yield differences in the ratio of larger aggregates (4-12 oligomers, or multimers) vs smaller aggregates (dimers) and monomers. For instance, FA on PLL displayed more monomers compared to glass and gelatine, which showed similar levels. FA on gelatine had larger clusters of paxillin, as indicated by the number of multimers. In a similar fashion as with RICS, a closer N&B analysis of regions with a size similar to the FAs resolved interesting differences. The apparent brightness frequency histograms at the peripheral FA were displaced towards higher values compared to central FA (frequency histograms are shifted towards the right in the X-axis, Fig 1. column 5). This effect is less remarkable when FAs are in contact with glass. A closer look at the oligomerisation distribution indicated that, whereas peripheral FAs on glass and PLL showed a higher ratio of monomers and dimers, central FAs were mostly composed by larger clusters (Fig. 1, column 5). Overall, peripheral FAs contained more multimers in contact with gelatin and PLL than on glass.

From this data we conclude that RICS and N&B combined sensitivity allows identifying different dynamics between cells attached to glass, PLL and gelatin. Paxillin diffuses faster at FAs formed on PLL or gelatin than those on glass. More importantly, these techniques provide information of differential dynamics in distinct but not-far-apart areas of the cell (peripheral and central FAs). Additionally, N&B can be used with the same acquisition parameters of RICS and supports the differences seen in diffusion among peripheral and central FAs by providing information of dissimilar aggregation of paxillin in these areas. Overall, we observed that slower adhesions show higher oligomerisation values as described by Digman et al. (7).

### 3.2 Dissecting the spatiotemporal dynamics of paxillin at specific self-assembly stages over the observed FA lifespan.

RICS analyses showed clear differences in diffusion between glass and PLL/gelatin. Further N&B observations inferred that FAs of cells on PLL and gelatine do not behave identically. The average intensity trace extracted from ≈7-10 min acquisitions (Fig. 1 column 3) (of either full-frame images or ROIs) showed abrupt changes in the fluorescence intensity at specific time points. These variations occurred several times along the acquisition and seemed cyclic, which might indicate that FAs were undergoing different assembly and disassembly stages. The periodic nature of these events lead us to hypothesise that some of the inconsistencies obtained before could be the result of masking particular events during long acquisitions. To confirm this hypothesis, we investigated the resolution capacity of the technique in shorter acquisition times (≈3 min). We re-analysed those groups of images that showed similar intensity trace changes between samples, specifically, those that exhibited a persistent increase in fluorescence intensity, which probably corresponded to the self-assembly of paxillin molecules at the FA (7, 29, 30). In Fig. 2 we show that RICS can discern discrete stages during the acquisition time, and resolve the spatiotemporal dynamics of paxillin at FAs of cells in contact with different surfaces. Consistent with the results observed in Fig. 1, overall paxillin diffusion was slower in cells on glass than on PLL or gelatine (Fig 2 column 2). We also confirmed that the molecular mobility of paxillin was consistently faster in central FAs than in peripheral FAs throughout all surfaces. During the assembly process (Fig. 2, right panel), peripheral FAs did not display large differences in paxillin diffusion between glass, PLL and gelatin. In contrast, the mobility at central FAs showed a clear trend: Gelatine>PLL>Glass. Interestingly, paxillin diffusion was faster during the assembly process (shorter time scale) in comparison to its mobility over the full-length acquisition, which involved several discrete assembly and disassembly processes (Fig. 2, columns 2 and 6).

**Figure 2:**
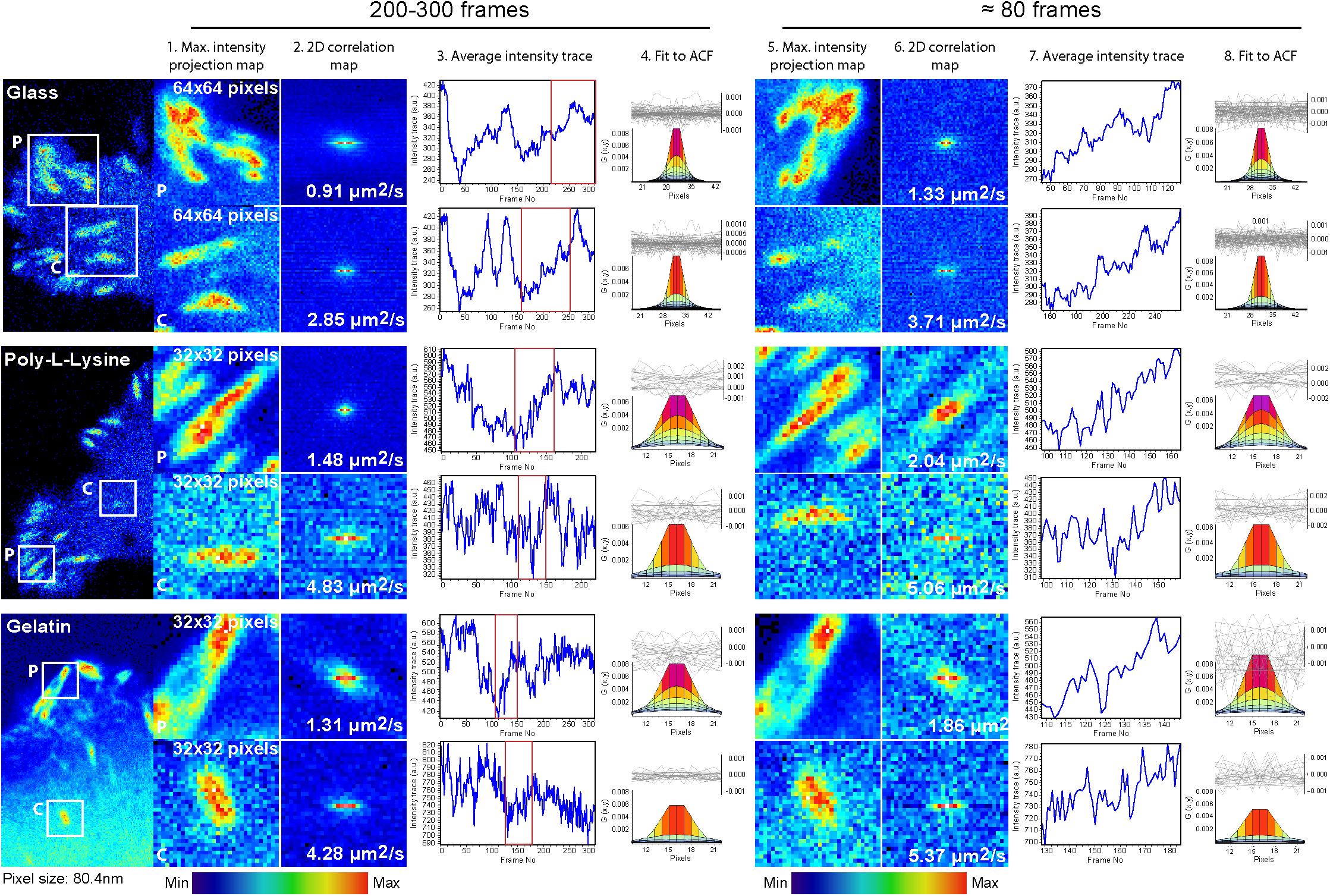
Optimisation of RICS analysis to characterise dynamic behaviour of Paxillin-EGFP diffusion at focal adhesions. MDA-MB-231 cells grown for 24h on glass, Poly-L-Lysine or gelatin, and imaged at 37<u>°</u>C and 5% CO_2_. RICS analyses were performed for paxillin-EGFP at peripheral FA (P) and at central FA (C). Analyses were done over 200-300 frames (7-10 min, left panel) or for about 80 frames (3 min, right panel). Representative maximum intensity projection maps (1 and 5), average intensity trace (2 and 6), 2D correlation maps and diffusion coefficient (3 and 7), and fit to ACF (4 and 8) resultant of RICS analysis are shown for each condition. Pixel size and colour-scale are shown below.

Previous studies using RICS to examine diffusion of cellular proteins, advise acquiring a number of frames in the range of ≈ 10-200 (28, 29). However, due to the stability of the z-focus and the fluorescence signal, as well as the viability of the cells during the acquisition, we acquired up to 200 to 300 frames. A longer period helped us to better understand the macromolecular context in which molecular dynamics of paxillin were taking place. As a result, we observed that in a time frame of ≈7 to 10 min, there were clear changes in all conditions in the fluorescence intensity along time (as shown in the average intensity trace in Fig.2, column 3), and these did not seem a consequence of drift in z or bleaching. These noticeable changes in intensity could correlate to different stages of development of the FA and, if that being the case, long acquisitions were probably masking discrete events leading to a loss in time-resolution. When we analysed shorter acquisition times (≈3 min) corresponding to paxillin molecular assembly, we did find different values of diffusion for every condition, all values corresponded with faster diffusion than those obtained in the analyses of longer acquisitions. Therefore, we conclude that RICS is capable of discriminating differential self-assembly molecular events of paxillin at different regions of the cell.

### 3.3 RICS and N&B analyses resolve the molecular spatiotemporal dynamics of intermediate stages during the observed FA lifespan

In different cell regions upon contact with diverse surfaces, RICS discriminates between paxillin dynamics at FAs. However, it is unclear whether RICS can detect differences between PLL and gelatine (that show a very similar behaviour). For this purpose, we focused only on cells in contact with PLL and gelatine. In Fig. 3 panel A, we see that overall (full time-frame acquisition) regardless of the contacting surface, the peripheral region have slower molecular mobility at the FA than the central side of the cell. On the contrary, central FAs displayed similar diffusion values, or slightly faster mobility in PLL.

**Figure 3:**
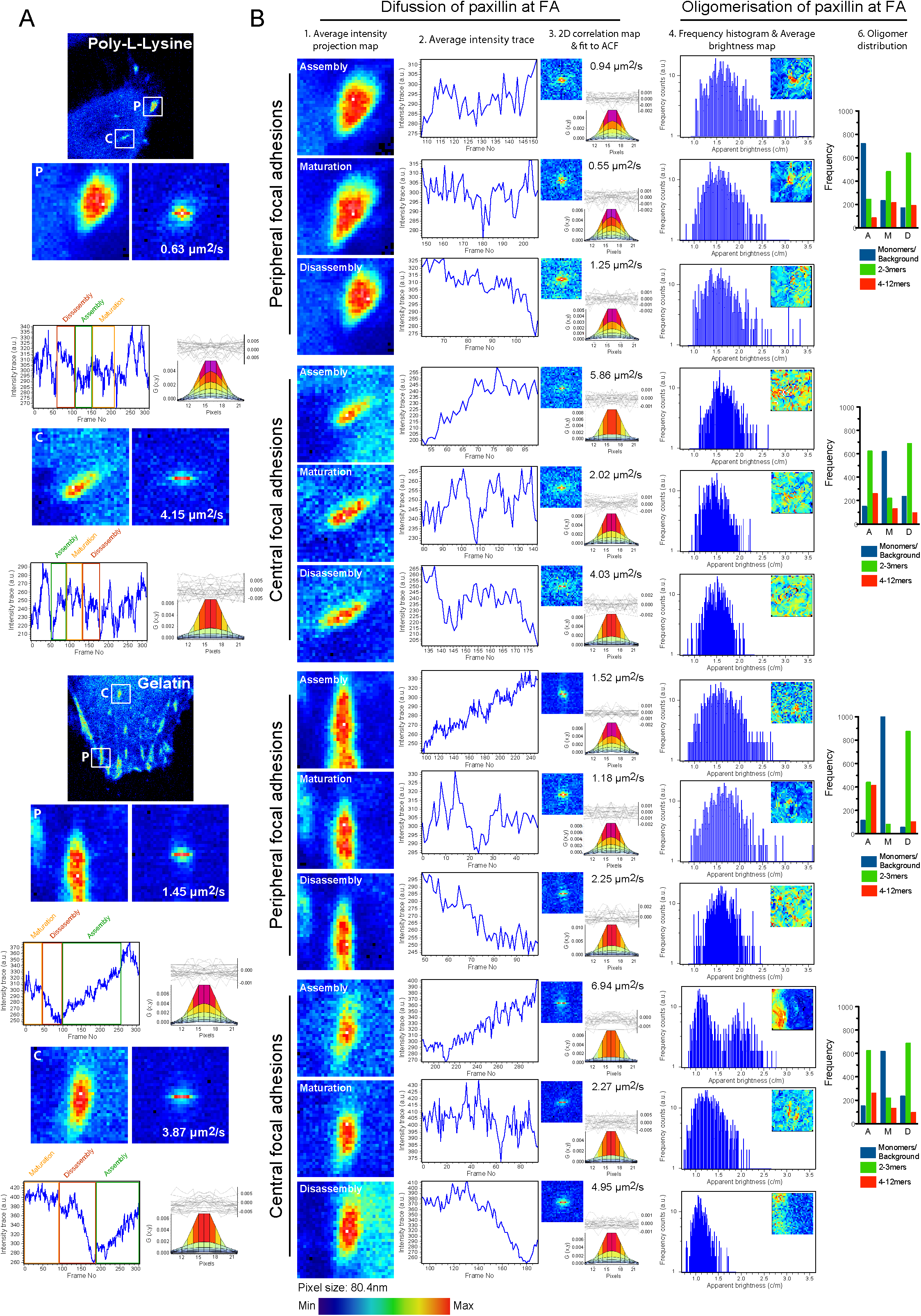
Characterisation molecular dynamics of different focal adhesion stages by RICS and N&B. MDA-MB-231 cells grown for 24h on Poly-L-Lysine (upper panel) or gelatin (lower panel) and imaged at 37<u>°</u>C and 5% CO_2_. (A) RICS analysis was performed for paxillin-EGFP at peripheral FA (P) and at central FA (C). (B) RICS and N&B analyses of FA assembly (obtained from the green perpendicular box depicted at the average intensity trace in panel A; characterised by prolonged/significant increase of fluorescence intensity as shown on intensity trace plots), FA maturation (obtained from the orange perpendicular box depicted at the average intensity trace in panel A; characterised by stages of stable fluorescence intensity) and FA disassembly (obtained from the red perpendicular box depicted at the average intensity trace in panel A; characterised by prolonged/significant reduction of fluorescence intensity). Representative maximum intensity projection maps (1), average intensity trace (2), 2D correlation and fit to ACF (3) resultant of RICS analysis are shown for the mention stages and conditions. Representative N&B analyses were also performed in assembly, maturation and disassembly stages. Frequency histogram (4) and distribution of oligomers (5) are shown for each condition. Pixel size and colour-scale are shown below.

As observed in Fig. 2, the average fluctuation of the intensity over time indicates again that several intermediate-self-assembly events may occur over the full acquisition time (Fig 3, panel A and B at the left hand side). These fluctuation traces suggest that during a time span of 7 to 10 min, FAs undergo different assembly and disassembly events, involving in some cases a maturation process in between. To identify discrete stages within the FA, we analysed three macromolecular events: assembly, corresponding to a sustained increase in intensity; maturation, defined by subtle changes in intensity and overall stability of the fluorescent signal; and disassembly, characterised by a persistent decrease in the intensity. To avoid misidentifying “disassembly” events as photo-bleaching phenomena, we excluded any sustained drop in signal intensity during the first 10-20 frames of the acquisition. These events could be resolved only analysing shorter time scales (maximum 3 min). To test the accuracy and robustness of these methods in short time scales, we again correlated quantitative information retrieved from the application of RICS and N&B to the development of FAs at the cell periphery and central area on cells on PLL or gelatine. Overall, during every single self-assembly stage (Assembly, A; Maturation, M; Disassembly, D) the diffusion of paxillin at FAs was faster on gelatine than on PLL (Fig. 3 B, column 3). This result reveals the high molecular spatiotemporal sensitivity of the method; both at regions with similar size to the actual FAs and even in smaller time scales, closer to the actual lifetime of the self-assembly undergoing at the FAs.

These findings are more evident if we observe the results of each individual surface separately. On PLL, peripheral FAs showed faster diffusion values during disassembly than throughout maturation, and we observed the slowest diffusion during the assembly (D>M>A). This trend changed at central FAs, in which assembly dynamics were faster than disassembly and maturation (A>D>M). On gelatine, the changes in diffusion values of peripheral FAs were as follows: D>A>M; whereas central FAs showed the same trend described for PLL: A>D>M. This may indicate that the peripheral FAs are more sensitive to the surfaces in contact, than the central ones. FAs may employ faster dynamics (turnover processes) for sensing purposes. In fact, peripheral FAs in PLL and gelatin have faster diffusion during disassembly (at least twice times faster than the diffusion coefficient during FA maturation) as shown by the 2D correlation maps and the diffusion coefficients in Fig. 3 B (column 3).

When we focused on the N&B analysis, we could appreciate changes in the aggregation of paxillin for different conditions. Overall, the assembly processes showed higher apparent brightness values than the maturation and disassembly processes (Fig 3, column 4). This could indicate that during the FA assembly, paxillin forms clusters via a higher degree of oligomerisation. This was especially noticeable in the case of gelatine, where for both, central and peripheral FAs mobilised higher paxillin oligomeric states (multimers), which was more evident during the assembly process. However, when we focused on the oligomerisation distribution analyses the data were inconclusive and therefore we were unable to withdraw any conclusions from the quantification of the oligomeric distribution in this set of experiments.

Based on these results, we can confirm that RICS is a robust tool for the study of paxillin dynamics at FA in live cells, being highly sensitive to diverse conditions: adhesion substrate, cellular localisation of the FAs and individual stages of self-assembly (assembly, disassembly and stabilisation or maturation within the structure of interest). A downside though was that N&B analyses did not provide enough evidence that would complement the RICS data; either because there were not significant differences in aggregation between assembly and disassembly, or due to the experimental conditions used. N&B was not sensitive enough to detect changes in aggregation, but is worth mentioning that we could correlate RICS and N&B analysis for the set of experiments shown in Figure 1. The main difference in the experiments shown in Fig. 1 and 3 is the number of frames analysed, since any other acquisition parameter did not change. This suggests what other studies have claimed (18, 19), that N&B requires a minimum number of frames in order to give accurate values, and that this technique is much more sensitive to different acquisition and detection settings. A conflict arises then in our study, since our results suggest that assembly and disassembly phenomena occur in a short time period (about 3 min in PLL and gelatine), and analysis including more frames would hide these processes. However, it is surprising that the apparent brightness histograms indicated differences at each of the individual paxillin self-assembly processes. A plausible hypothesis is that the oligomerisation process is much more sensitive to the stage of cell activation at the time of the acquisition, which will depend on intrinsic and extrinsic cues. Our study so far has investigated and compared the diffusion and clustering dynamics of paxillin at FAs in a heterogeneous population of cells. To overcome this plausible limitation, for this purpose though, it may be advantageous to investigate all these processes at the single-cell level.

### 3.4 RICS-N&B combined analysis resolves the molecular spatiotemporal dynamics and clustering of intermediate stages and identifies a polarised behaviour in the dynamics of FA-associated paxillin

To discard that the previous inconclusive N&B results were due to high levels of cell heterogeneity (such as different states of activation, migratory capabilities, etc), we employed single-cell approaches. We focused on gelatine-coated surfaces, first because is physiologically more relevant than glass or PLL coating, and second because cells on gelatin displayed the fastest dynamics and thus we could challenge the sensitivity of these techniques. In Fig. 4 we show a migrating cell that clearly showed three different areas containing FAs: a distal or peripheral area with an active lamellipodium; an intermediate or peripheral-central area that contained FAs in the periphery but further away from the lamellipodium; and finally a central area. We selected two ROIs per area that contained FAs and performed RICS and N&B analyses, this time focusing only in the assembly and disassembly stages. We did not investigate the stabilisation or maturation process since previous analyses of oligomerisation distribution indicated this process was widely heterogeneous.

**Figure 4:**
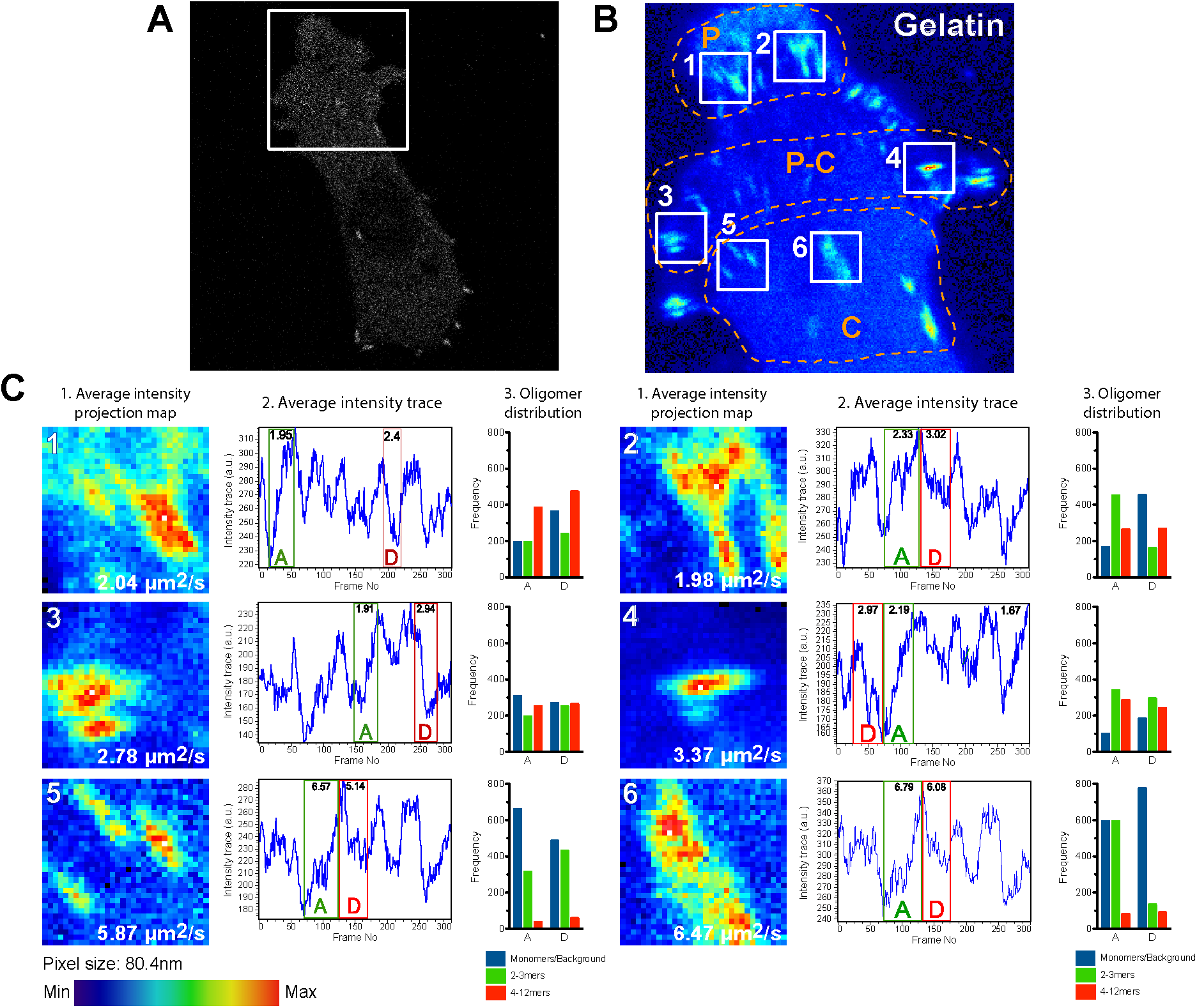
Diffusion of FA-associated paxillin is polarised alongside the central-peripheral axis. MDA-MB-231 cells grown for 24h on gelatin and imaged at 37<u>°</u>C and 5% CO_2_. (A) Confocal image of paxillin-EGFP. (B) Average intensity projection map of inset shown in A. Dashed line limits the different analysed areas. (C) RICS and N&B analyses performed on ROIs shown in B for paxillin-EGFP at peripheral FA (P), peripheral-central FA (P-C) and at central FA (C). Maximum intensity projection maps and diffusion coefficient (1), average intensity trace (2) and distribution of oligomers (3) are shown for each condition. Pixel size and colour-scale are shown below.

At the single-cell level, paxillin dynamics of FAs on gelatine were similar to those shown in Fig. 3. Interestingly, we observed a gradual increase in the diffusion values from the cell periphery towards the centre of the cell (Fig. 4 C, column 1), and opposite to the direction of the cell movement. Moreover, we observed a similar trend when we focused only on assembly and disassembly processes (Fig. 4 C, column 2). For instance, at peripheral and central-peripheral FAs the assembly was slower than the disassembly, while at central FAs the assembly was faster than the disassembly (Fig. 4 C, column 2).

The distribution of paxillin oligomers in FA on gelatine also showed a polarised behaviour that correlated with the direction of movement. We observed more molecules of low oligomerisation state the central part of the cell that gradually decreased towards the leading edge. Accordingly, the distribution of molecules of higher oligomeric state was the exact opposite, with a much higher number of multimers at the leading edge. This indicates a reversed flow of molecular diffusion and at the FAs between low and high oligomeric states. We observed a similar distribution in gelatine (though not so detailed) in Fig. 1. In addition to the differences observed between areas within the cell, we also detected changes in oligomerisation among assembly and disassembly events in peripheral and central FAs. At peripheral FAs, we detected less molecules of low degree of oligomerisation than molecules of high oligomerisation degree (multimers) during the assembly process. However, when FAs transitioned from assembly to disassembly, we observed about a 40% decrease of dimers, an increase in monomers (more than twice), and a small increase in multimers (15% more). At central-peripheral FAs though, the levels of monomers, dimers and multimers seemed fairly constant during the full assembly/disassembly process. Finally, at central FAs, where paxillin diffusion is the fastest, we observed that the percentage of dimers decreased almost a 40%, whereas monomers and multimers levels increased about 25% each.

Previous results from Digman et al. also support the likely synchronisation in dynamics of paxillin in particular areas of the cell. However, our results indicate dissimilar oligomerisation states during assembly and disassembly events (7). These differences can reside in the cell type and substrate used in each study (as we could observed in figure 1 and 3), or how cells respond to chemical signals from the growth medium. In addition, in regards to the oligomerisation values obtained by N&B, we noted that quantification using frequency analyses weights more the changes in the numbers of monomers and dimers than those of multimers, obscuring the relative changes between the conditions tested.

These results demonstrate that the molecular mobility of paxillin and its oligomerisation during assembly/disassembly events at FA can be resolved combining the analysis of RICS and N&B at the single cell level. This opens a huge potential for resolving spatiotemporal dynamics that occur at different regions of the cells and with different speed. The information extracted at the single-cell level could then be extrapolated to larger populations by increasing the number of studied cells. A problem that might arise though is the putative difficulty of finding a value or characteristic to normalise the different cells, and thus be able to compare among different conditions. To start with, in some cases it might be better off, investigating the dynamics and oligomerisation distribution in separate experiments employing different acquisition conditions. Then, with this *a priori* knowledge in hand, one could increase the level of detail by investigating single cell events, in a similar fashion as it has been done in this report. Definitively, the level of detail and information that can be retrieved and combined by correlating both techniques makes this approach extremely tempting.

## Conclusions

In this work we have exhaustively examined the sensitivity and limitations of RICS and N&B as methods to identify and characterise the molecular dynamics of paxillin-EGFP at the FAs. For this study, we employed MDA-MB-231 epithelial cells. These cells, isolated from a highly invasive breast carcinoma, display an elevated migratory and invasive behaviour that is highly dependent on the surrounding (34). To study the FAs of these cells at the molecular level, we used three different attachment conditions: non-coated glass, poly-L-lysine (PLL) and gelatine. Among these substrate conditions, only the attachment to gelatine is mediated by integrins (35), whereas cell attachment to glass and PLL occurs via interaction between the polyanions from the cell surface and polycations in the glass or PLL coating. Based on these mechanisms of cell-surface interaction and previously published data (32, 33) seemed plausible that these cells would attach more stably to glass and PLL, and would be more motile and dynamic on gelatine. Initial observations of the paxillin diffusion at FAs upon contact with these surfaces indicated that cells on glass had a slower molecular diffusion than those on PLL or gelatin, suggesting a lower interchange of molecules at the FA and therefore a more stable adhesion. Former works have characterised the molecular dynamics of paxillin upon stimulation with substrates have been performed using fibronectin (7, 30), but studies regarding cell attachment to different substrates do support our observations (32, 33). Several reports have addressed the heterogeneity of the dynamic behaviour of FAs (5-7, 30), however our observations suggest that in spite of existing a great variability regarding the diffusion of paxillin molecules and specially the oligomerisation processes at FAs, the appropriate approach can lead to noteworthy results. For instance, we observe that RICS detects differential adhesion dynamics on cells attached on glass, PLL or gelatin, comprising slower paxillin diffusion at FAs on glass and faster diffusion on gelatine. In addition, the spatiotemporal resolution of RICS allows the detection and quantification of differential assembly and disassembly dynamics at FAs from different cellular regions and upon contact to different surfaces. Employing simultaneously RICS and N&B allows gaining further insights on the molecular mechanisms of paxillin diffusion and aggregation at FAs in the aforementioned conditions in particular at the single-cell level. In these conditions, we are able to resolve the molecular spatiotemporal dynamics and clustering of intermediate stages and identify a polarised behaviour in the dynamics of FA-associated paxillin. We saw higher number of molecules with low degree of oligomerisation at the central part of the cell, which gradually decreased towards the leading edge. Coordinatelly, the distribution of molecules with higher degree of oligomerisation was the exact opposite, with a much higher number of multimers at the leading edge. Overall, this suggests a reversed flow of molecular diffusion and at the FAs between low and high oligomeric states.

Although most studies have tackled the issue of the high variability in FA dynamics by studying specific stages of the FA development such as nascent or mature/stable FA (7, 30, 36), these approaches rely on the correct identification of these stages or analysing a huge cell population. We propose a single-cell based approach and show that this methodology can distinguish differential dynamic behaviours among various subcellular localisations.

## Acknowledgements

We are grateful to the STFC (Science and Technology Facility Council) for application microscopy time. J.B.S. and E.G. are grateful for support from a Marie Curie Career Integration Grant “NanodynacTCELLvation” PCIG13- GA-2013-618914.

---

**Figure S1:**
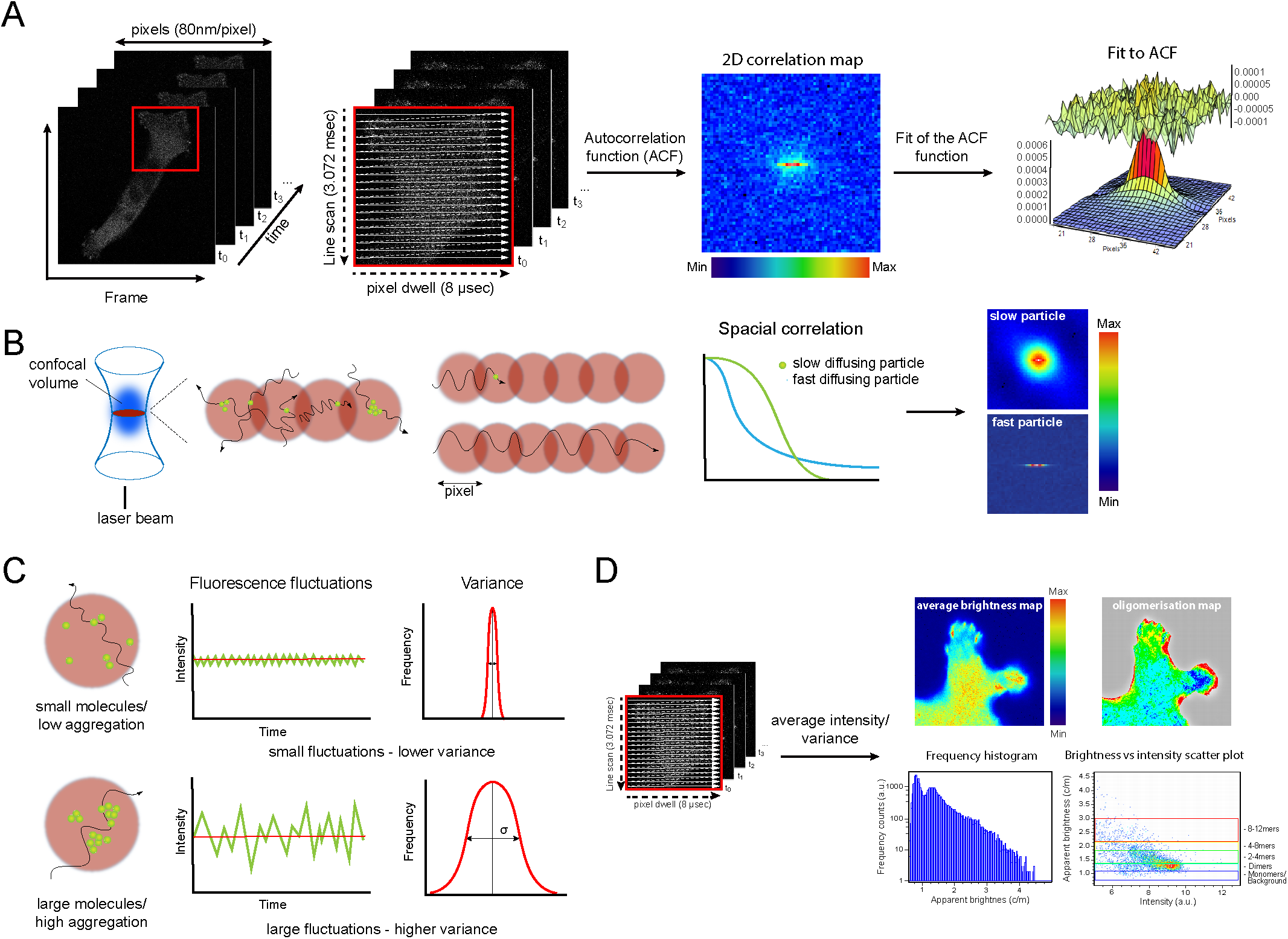
Description of RICS and N&B methods. (A) Schematic of image acquisition and analysis performed by RICS: series of 200-300 images of 256 pixels by 256 pixels (20.58μm by 20.58 μm) were continuously collected at a pixel dwell time of 8 μsec. These images were processed using SimFCS4 to calculate the autocorrelation function (ACF) and subsequently perform a fit to this ACF. From this fitting, we obtained a value of diffusion and a range of residual values. (B) Graphic exemplification of slow and fast particle diffusion during line scanning and correlation analyses. Point illumination originates a confocal volume or PSF through which fluorescent molecules cross at different speeds depending on their size, molecular interactions and steric hindrance. Slow molecules will correlate mostly correlate with neighbour pixels/confocal volumes whereas faster particles show higher correlation with pixels that are further away. (C) N&B methodology: representation of moment analysis based on determining the average intensity (first moment) and the variance (second moment) of the fluorescence intensity fluctuation. For two equal fluorescence intensities, a larger variance implies less contribution of these molecules to the average. (D) The raster analysis provides a map of number and brightness for every pixel in the image, from which frequency of monomers/background, dimers and bigger aggregates can be extracted.

**Figure S2:**
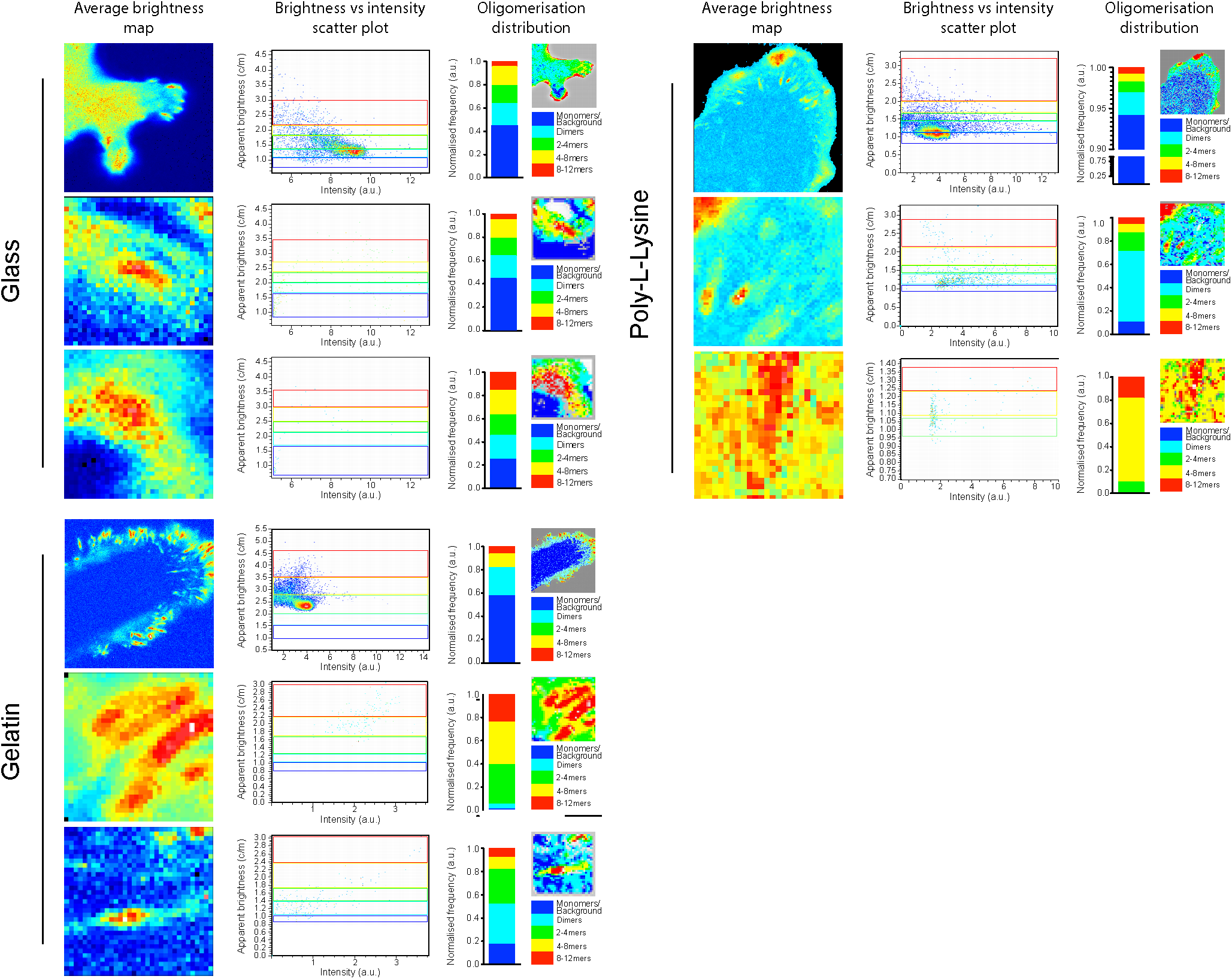
Extended N&B analysis of paxillin-EGFP oligomeric state at focal adhesions in cells grown on difference surfaces. MDA-MB-231 cells grown for 24h on glass, Poly-L-Lysine or gelatin, and imaged at 37<u>°</u>C and 5% CO_2_. N&B analysis were performed for total paxillin-EGFP (top), paxillin-EGFP at peripheral FA (medium) and at central FA (bottom). Analyses were performed over 200-300 frames (over 7-10 min). Representative average brightness maps, brightness vs intensity scatter plot, mirrored oligomerisation map (see material and methods section) and quantified oligomerisation distribution.

## References

1. Carragher NO, Frame MC. Focal adhesion and actin dynamics: a place where kinases and proteases meet to promote invasion. Trends Cell Biol. 2004;14(5):241-9.

2. Schiller HB, Fassler R. Mechanosensitivity and compositional dynamics of cell-matrix adhesions. EMBO Rep. 2013;14(6):509-19.

3. Humphrey JD, Dufresne ER, Schwartz MA. Mechanotransduction and extracellular matrix homeostasis. Nat Rev Mol Cell Biol. 2014;15(12):802-12.

4. Livne A, Geiger B. The inner workings of stress fibers - from contractile machinery to focal adhesions and back. J Cell Sci. 2016;129(7):1293-304.

5. Choi CK, Vicente-Manzanares M, Zareno J, Whitmore LA, Mogilner A, Horwitz AR. Actin and alpha-actinin orchestrate the assembly and maturation of nascent adhesions in a myosin II motor-independent manner. Nat Cell Biol. 2008;10(9):1039-50.

6. Shroff H, Galbraith CG, Galbraith JA, Betzig E. Live-cell photoactivated localization microscopy of nanoscale adhesion dynamics. Nat Methods. 2008;5(5):417-23.

7. Digman MA, Brown CM, Horwitz AR, Mantulin WW, Gratton E. Paxillin dynamics measured during adhesion assembly and disassembly by correlation spectroscopy. Biophys J. 2008;94(7):2819-31.

8. Brown CM, Dalal RB, Hebert B, Digman MA, Horwitz AR, Gratton E. Raster image correlation spectroscopy (RICS) for measuring fast protein dynamics and concentrations with a commercial laser scanning confocal microscope. J Microsc. 2008;229(Pt 1):78-91.

9. Jacobson K, Derzko Z, Wu ES, Hou Y, Poste G. Measurement of the lateral mobility of cell surface components in single, living cells by fluorescence recovery after photobleaching. J Supramol Struct. 1976;5(4):565(417)–576(428).

10. Magde D, Webb WW, Elson E. Thermodynamic Fluctuations in a Reacting System-Measurement by Fluorescence Correlation Spectroscopy. Phys Rev Lett. 1972;29(11):705-&.

11. Digman MA, Brown CM, Sengupta P, Wiseman PW, Horwitz AR, Gratton E. Measuring fast dynamics in solutions and cells with a laser scanning microscope. Biophys J. 2005;89(2):1317-27.

12. Qian H, Sheetz MP, Elson EL. Single particle tracking. Analysis of diffusion and flow in two-dimensional systems. Biophys J. 1991;60(4):910-21.

13. Clausen MP, Sezgin E, Bernardino de la Serna J, Waithe D, Lagerholm BC, Eggeling C. A straightforward approach for gated STED-FCS to investigate lipid membrane dynamics. Methods. 2015;88:67-75.

14. Eggeling C, Ringemann C, Medda R, Schwarzmann G, Sandhoff K, Polyakova S, et al. Direct observation of the nanoscale dynamics of membrane lipids in a living cell. Nature. 2009;457(7233):1159-62.

15. Hedde PN, Dorlich RM, Blomley R, Gradl D, Oppong E, Cato AC, et al. Stimulated emission depletion-based raster image correlation spectroscopy reveals biomolecular dynamics in live cells. Nat Commun. 2013;4:2093.

16. Bernardino de la Serna J, Schutz GJ, Eggeling C, Cebecauer M. There Is No Simple Model of the Plasma Membrane Organization. Front Cell Dev Biol. 2016;4:106.

17. Qian H, Elson EL. Distribution of molecular aggregation by analysis of fluctuation moments. Proc Natl Acad Sci U S A. 1990;87(14):5479-83.

18. Dalal RB, Digman MA, Horwitz AF, Vetri V, Gratton E. Determination of particle number and brightness using a laser scanning confocal microscope operating in the analog mode. Microsc Res Tech. 2008;71(1):69-81.

19. Digman MA, Dalal R, Horwitz AF, Gratton E. Mapping the number of molecules and brightness in the laser scanning microscope. Biophys J. 2008;94(6):2320-32.

20. Youker RT, Teng H. Measuring protein dynamics in live cells: protocols and practical considerations for fluorescence fluctuation microscopy. J Biomed Opt. 2014;19(9):90801.

21. Gielen E, Smisdom N, De Clercq B, Vandeven M, Gijsbers R, Debyser Z, et al. Diffusion of myelin oligodendrocyte glycoprotein in living OLN-93 cells investigated by raster-scanning image correlation spectroscopy (RICS). J Fluoresc. 2008;18(5):813-9.

22. Vendelin M, Birkedal R. Anisotropic diffusion of fluorescently labeled ATP in rat cardiomyocytes determined by raster image correlation spectroscopy. Am J Physiol Cell Physiol. 2008;295(5):C1302-15.

23. Sasaki A, Yamamoto J, Jin T, Kinjo M. Raster image cross-correlation analysis for spatiotemporal visualization of intracellular degradation activities against exogenous DNAs. Sci Rep. 2015;5:14428.

24. Jones DM, Alvarez LA, Nolan R, Ferriz M, Sainz Urruela R, Massana-Munoz X, et al. Dynamin-2 Stabilizes the HIV-1 Fusion Pore with a Low Oligomeric State. Cell Rep. 2017;18(2):443-53.

25. Hellriegel C, Caiolfa VR, Corti V, Sidenius N, Zamai M. Number and brightness image analysis reveals ATF-induced dimerization kinetics of uPAR in the cell membrane. FASEB J. 2011;25(9):2883-97.

26. Nolan R, Alvarez LAJ, Elegheert J, Iliopoulou M, Jakobsdottir GM, Rodriguez-Munoz M, et al. nandb-number and brightness in R with a novel automatic detrending algorithm. Bioinformatics. 2017;33(21):3508-10.

27. Liang EI, Mah EJ, Yee AF, Digman MA. Correlation of focal adhesion assembly and disassembly with cell migration on nanotopography. Integr Biol (Camb). 2017;9(2):145-55.

28. Rossow MJ, Sasaki JM, Digman MA, Gratton E. Raster image correlation spectroscopy in live cells. Nat Protoc. 2010;5(11):1761-74.

29. Digman MA, Wiseman PW, Horwitz AR, Gratton E. Detecting protein complexes in living cells from laser scanning confocal image sequences by the cross correlation raster image spectroscopy method. Biophys J. 2009;96(2):707-16.

30. Shibata AC, Fujiwara TK, Chen L, Suzuki KG, Ishikawa Y, Nemoto YL, et al. Archipelago architecture of the focal adhesion: membrane molecules freely enter and exit from the focal adhesion zone. Cytoskeleton (Hoboken). 2012;69(6):380-92.

31. Wilson CJ, Clegg RE, Leavesley DI, Pearcy MJ. Mediation of biomaterial-cell interactions by adsorbed proteins: a review. Tissue Eng. 2005;11(1-2):1-18.

32. Davidenko N, Schuster CF, Bax DV, Farndale RW, Hamaia S, Best SM, et al. Evaluation of cell binding to collagen and gelatin: a study of the effect of 2D and 3D architecture and surface chemistry. J Mater Sci Mater Med. 2016;27(10):148.

33. Liberio MS, Sadowski MC, Soekmadji C, Davis RA, Nelson CC. Differential effects of tissue culture coating substrates on prostate cancer cell adherence, morphology and behavior. PLoS One. 2014;9(11):e112122.

34. Garcia E, Ragazzini C, Yu X, Cuesta-Garcia E, Bernardino de la Serna J, Zech T, et al. WIP and WICH/WIRE co-ordinately control invadopodium formation and maturation in human breast cancer cell invasion. Sci Rep. 2016;6:23590.

35. Costa P, Scales TM, Ivaska J, Parsons M. Integrin-specific control of focal adhesion kinase and RhoA regulates membrane protrusion and invasion. PLoS One. 2013;8(9):e74659.

36. Kaushik S, Ravi A, Hameed FM, Low BC. Concerted Modulation of Paxillin Dynamics at Focal Adhesions by Deleted in Liver Cancer-1 and Focal Adhesion Kinase During Early Cell Spreading. Cytoskeleton. 2014;71(12):677-94.

